# Viable mutants of essential genes in *Physcomitrium patens* as tools for studying primary metabolic processes

**DOI:** 10.1101/2023.09.20.558677

**Authors:** Tegan M. Haslam, Cornelia Herrfurth, Ivo Feussner

**Affiliations:** University of Goettingen, Albrecht-von-Haller-Institute for Plant Sciences, Dept. of Plant Biochemistry, D-37077 Goettingen, Germany; University of Goettingen, Goettingen Center for Molecular Biosciences (GZMB), Service Unit for Metabolomics and Lipidomics, D-37077 Goettingen, Germany; University of Goettingen, Goettingen Center for Molecular Biosciences (GZMB), Dept. of Plant Biochemistry, D-37077 Goettingen, Germany

## Abstract

Sphingolipids are essential components of plant cells, which have been notoriously difficult to study in part due to pleiotropic or lethal knock-out mutant phenotypes. By relying on alternative end-joining of double stranded breaks, we successfully used CRISPR/Cas9 mutagenesis to generate a population of diverse, viable mutant alleles of genes required for sphingolipid assembly in totipotent protoplasts of the moss *Physcomitrium patens*. We targeted the *INOSITOL PHOSPHORYLCERAMIDE SYNTHASE* (*IPCS*) gene family, which catalyzes the committed step in the synthesis of glycosyl inositol phosphorylceramides (GIPCs), the most abundant class of sphingolipids found in plants. We isolated knock-out single mutants and knock-down higher-order mutants showing a spectrum of deficiencies in GIPC content. Remarkably, we also identified two mutant alleles accumulating inositol phosphorylceramides, the direct products of IPCS activity, and provide our best explanation for this unexpected phenotype. Our approach is broadly applicable for studying essential genes and gene families, and for obtaining unusual lesions within a gene of interest.

## Introduction

Glycosyl inositol phosphorylceramides (GIPCs) are essential lipids in plants. They make up approximately one third of the plasma membrane lipid content (Bahammou et al., 2023), and are specifically enriched in its outer leaflet (Tjellström et al., 2010). This localization and enrichment fit with the demonstrated roles of GIPCs in signal transduction between the apoplast and symplast (Jiang et al., 2019), interactions with both pathogenic and symbiotic microorganisms (Lenarčič et al., 2017; Moore et al., 2021), and influence over the electrochemical polarization and physical organization of the plasma membrane (Mamode Cassim et al., 2020).

GIPCs consist of a ceramide backbone that contains a saturated or monounsaturated very-long-chain fatty acid (VLCFA) and a poly-hydroxylated long-chain-base (LCB) moiety with a polar sugar headgroup (reviewed in Luttgeharm et al., 2016; Haslam and Feussner, 2022). The ceramide structure supports increased conformational order in membranes or domains of membranes where GIPCs accumulate, as their poly-hydroxylated acyl and LCB tails interact via hydrogen bonds with phytosterols that are also characteristic of ordered lipid membranes (Lingwood and Simons, 2010; Cassim et al., 2021). The presence of VLCFA-containing GIPCs is associated with increased membrane thickness in both plants (Cassim et al., 2021) and in the yeast *Saccharomyces cerevisiae*, where analogous inositol phosphorylceramides (IPCs), mannosylated IPCs (MIPCs), and mannosylated diinositol phosphorylceramides (M(IP)_2_Cs) accumulate (Levine et al., 2000). VLCFA moieties of GIPCs have also been found to be directly required for polar sorting of PIN2 auxin efflux carriers mediated by phosphoinositides (Wattelet-Boyer et al., 2016; Ito et al., 2021).

The defining feature of GIPCs is their hexosyl-glucuronic acid-inositol-phosphate headgroup attached to position 1 of the LCB, which protrudes into the apoplast. The first committed step in GIPC synthesis is the addition of the first moiety of the headgroup, inositol phosphate, which is transferred from phosphatidyl inositol onto ceramide by INOSITOL PHOSPHATE CERAMIDE SYNTHASE (IPCS) (Wang et al., 2008) (Figure 1A). This step is common to plant GIPCs and fungal and protozoan IPCs/MIPCs, and the responsible enzymes have sequence similarity to enzymes transferring phosphocholine residues to ceramides to produce sphingomyelin in animals (Denny et al., 2007; Mina et al., 2010). Next, a glucuronic acid residue is added by INOSITOL PHOSPHATE GLUCURONOSYL TRANSFERASE (Rennie et al., 2014; Tartaglio et al., 2017). This step is specific to GIPC assembly, and confers a negative charge to the headgroup that contributes to plasma membrane electric potential (Cassim et al., 2021). Recent evidence suggests that the negatively charged sugar acid interacts with cations in the apoplast, and thereby senses and directs physiological responses to environmental conditions (Jiang et al., 2019). Beyond the glucuronic acid residue, GIPC headgroups may contain anywhere from a single sugar moiety up to 19-20, based on the ratio of carbohydrate to lipid in purified extracts (Kaul and Lester, 1975). Generally, these additional sugar moieties are classified as hexose (Hex) or N-acetyl hexosamine (HexNAc). Whether the final products include one or twenty, Hex or HexNAc, residues, they are generally collectively referred to as GIPCs except in instances where the headgroups are being specifically identified and distinguished. Most studies measure only mono- and di-hexosylated GIPCs, due to technical challenges in measuring increasingly polar, high molecular weight lipids. The chemical composition of the headgroup varies among plant taxa (Cacas et al., 2013), and within a given plant between different tissue types (Luttgeharm et al., 2015; Tellier et al., 2014). Specific headgroups can be recognized and are essential for interactions with microbial symbionts (Moore et al., 2021) and pathogens (Lenarčič et al., 2017).

**Figure 1:**
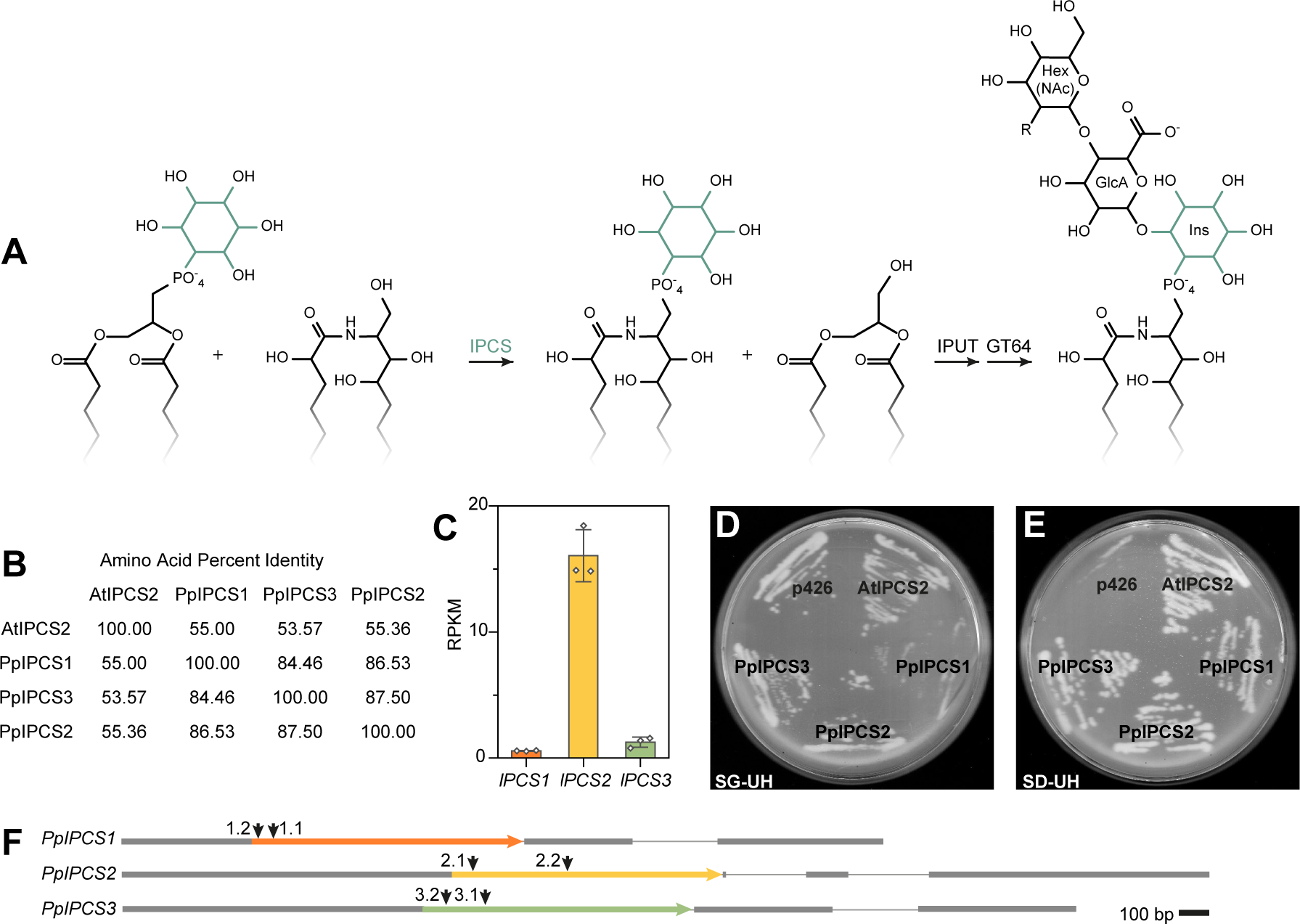
Identification of candidate *INOSITOL PHOSPHORYLCERAMIDE SYNTHASE* (*IPCS*) genes in *Physcomitrium*. **A** Assembly of a typical GIPC headgroup. The committed step is the transfer of inositol phosphate from phosphatidyl inositol onto a ceramide backbone, catalyzed by IPCS, producing inositol phosphoryl ceramide and diacylglycerol. **B** Percent identity matrix of *Arabidopsis* IPCS2 and candidate *Physcomitrium* IPCS proteins. **C** Expression profile of *PpIPCS* genes, from mature gametophores. Values are averages of three biological replicates, error bars represent standard deviation. **D & E** Complementation of *S. cerevisiae* YPH499-HIS-GAL1::AUR1 conditional mutant with AtIPCS2 (positive control), PpIPCS1, PpIPCS2, and PpIPCS3, with the empty p426 vector as a negative control. Endogenous *AUR1* is expressed in cells grown on galactose-containing media (SG-UH, **D**), and suppressed in cells grown on media lacking galactose, containing glucose (SD-UH, **E**). **F** Schematic of *PpIPCS* genes, with coding sequence as colored arrows, UTRs as grey bars, and intronic sequence as a fine grey line. Positions of CRISPR targets are marked with arrows (↓).

We have learned much about the synthesis and functions of GIPCs from genetic studies in *Arabidopsis thaliana*, *Oryza sativa*, and recently, *Medicago truncatula* (Mortimer and Scheller, 2020), which have described consequences of deficiencies in late steps in the assembly of the complex GIPC headgroups, such as *GIPC MANNOSYL TRANSFERASE* (*GMT*) (Fang et al., 2016) and *GLUCOSAMINE INOSITOL PHOSPHORYLCERAMIDE TRANSFERASE1* (*GINT1*) (Ishikawa et al., 2018; Moore et al., 2021), and the sugar transporters responsible for making substrate available in the Golgi for headgroup assembly, *GOLGI-LOCALIZED NUCLEOTIDE SUGAR TRANSPORTER1* (Mortimer et al., 2013) and *2* (Jing et al., 2021) (*GONST1*/2) and *UDP-GlcNAc TRANSPORTER1* (*UGNT1*) (Ebert et al., 2018). These have produced dramatic phenotypes and revealed a diverse set of functions for specific headgroup moieties. Work on mutants affected in the modification of ceramide backbones have also provided insight to how desaturations and hydroxylations of GIPCs affect membrane dynamics (Markham et al., 2011; Wattelet-Boyer et al., 2016).

Efforts to target early steps in GIPC assembly by direct, knock-out mutagenesis have been challenged by the essential nature of GIPCs and their myriad functions. The first step of GIPC headgroup assembly is the addition of inositol phosphate to ceramide catalyzed by INOSITOL PHOSPHORYLCERAMIDE SYNTHASE (Figure 1A). In *A. thaliana* three loci, *IPCS1, IPCS2/ERH1,* and *IPCS3* encode proteins with this catalytic activity (Wang et al., 2008; Mina et al., 2010). Single *ipcs2/erh1* mutants have a dwarf phenotype and develop spontaneous, hypersensitive-response-like cell death; however, though *ipcs2/erh1* plants accumulate ceramides and hydroxyceramides, they display no substantial decrease in GIPC content (Wang et al., 2008). Attempts to isolate *ipcs1 ipcs2 ipcs3* triple mutants have been unsuccessful, fitting with the expectation that such mutants should be lethal (Mina et al., 2010). *ipcs1 ipcs2* double mutants are severely stunted and seedling lethal (Ito et al., 2021). Therefore, *IPCS1 IPCS2* artificial micro RNA (amiRNA) lines were generated, which present a strong reduction in *IPCS* transcripts and defects in polar protein transport. However, the amiRNA lines displayed only modest reductions in GIPC levels, rendering the identification of additional functions of GIPCs difficult (Ito et al., 2021).

To improve our understanding of the biological functions of sphingolipids in plants, our group and others have used the model moss *Physcomitrium patens* (formerly *Physcomitrella patens*) (Medina et al., 2019) for its relatively simple developmental patterning and organ structure (Resemann et al., 2021; Gömann et al., 2021a, 2021b; Steinberger et al., 2021; Haslam and Feussner, 2022). Further, *P. patens* is a wonderful model for expanding the phylodiversity of our knowledge in plant biology (Rensing et al., 2020). We anticipated that *P. patens* could also be practically useful for studying essential genes and gene families, such as *IPCS*s, due to one option for CRISPR/Cas9 mutagenesis that is uniquely easy and efficient in *P. patens*.

Two basic strategies for CRISPR/Cas9 genome editing commonly used in plants are to (1) use two single guide RNAs (sgRNAs) targeting both ends of a coding sequence to produce a large deletion, and (2) use one sgRNA, and rely on mutations introduced by alternative end joining reconnecting the single Cas9-induced double-stranded break. This second approach will generate diverse lesions around the selected target (Collonnier et al., 2017; Lopez-Obando et al., 2016); we can expect knock-outs, a range of knock-down lesions, and perhaps also lesions that introduce changes in gene expression, gene product localization, or acquire modified or novel gene functions. Because knock-outs resulting from frame-shifts are the most probable outcome, acquiring more interesting alleles is primarily limited by the number of independently-mutagenized individuals produced for screening. As *P. patens* is one of few plant models that can be easily regenerated from single protoplasts, producing populations of independently-transformed, and independently-CRISPR-Cas9-mutagenized protoplasts, is uniquely efficient in this system.

We used this approach to mutagenize the *IPCS* gene family, and collected and analyzed a population of mutants with knock-out, knock-down, and modified functionality phenotypes. We anticipate this method will be broadly applicable to study essential gene functions, and to find unexpected and unique mutant alleles in a semi-targeted manner.

## Results

### The *P. patens* genome encodes three functional *IPCS* genes

Three candidate *IPCS* genes in the *P. patens* Gransden V3.3 genome were identified and their gene products compared (Supplemental Figure 1, Figure 1B). Two conserved motifs common to IPCSs (Denny et al., 2007; Wang et al., 2008) are present in all three sequences (Supplemental Figure 1). Expression profiles of the candidate genes were determined from RNA Seq data of wild-type Gransden 2004 gametophores. *PpIPCS2* is approximately 28X more strongly expressed than *PpIPCS1* and 13X more strongly expressed than *PpIPCS3* in mature gametophores (Figure 1C; Supplemental Table 1). Similar trends were found on PEATMoss, and the *Physcomitrella* eFP browser (Winter et al., 2007; Ortiz-Ramírez et al., 2016; Perroud et al., 2018; Fernandez-Pozo et al., 2020)(Supplemental Table 2).

To determine whether annotated *IPCS* genes function as IPCSs *in vivo*, the *IPCS1*, *IPCS2*, and *IPCS3* genes were cloned into a p426 expression vector (Mumberg et al., 1995; Sikorski and Hieter, 1989) for heterologous expression driven by the strong, constitutive *GPD3* promoter in baker’s yeast, *Saccharomyces cerevisiae*. A conditional mutant was used or this experiment, *YPH499–HIS3–GAL1:AUR1* (Denny et al., 2007), kindly provided by Prof. Dr. Ralph T. Schwarz and Dr. med vet. Hosam Shams-Eldin (Institute for Virology - BMFZ, Philipps-Universität Marburg). The promoter of the endogenous yeast *IPCS*, *AUR1*, is replaced by a galactose-inducible promoter in this strain; therefore, *AUR1* is expressed when cells are grown on galactose, and suppressed on glucose. Arabidopsis *IPCS2* was used as a positive control for this experiment (Mina et al., 2010). All three *IPCS*s rescued growth of the *YPH499-HIS3-GAL1:AUR1* strain (Figure 1D, E), indicating that in yeast cells the three can function as IPCSs.

As all three *PpIPCS* gene loci are functional IPCSs and are expressed in gametophytic tissues, it is likely that a triple mutant will be necessary to observe a loss-of-function phenotype.

### Single and higher-order mutants were generated by CRISPR/Cas9 mutagenesis

A series of *ipcs* single mutants was generated by CRISPR/Cas9 (Collonnier et al., 2017; Lopez-Obando et al., 2016), targeting the positions indicated in Figure 1F. Four independent alleles were selected for each gene locus for further study, which were expected to have true knock-out lesions (Supplemental Table 3). A subtle growth defect was observed in all *ipcs2* mutants (Figure 2), while *ipcs1* and *ipcs3* mutants were indistinguishable from the wild type.

**Figure 2:**
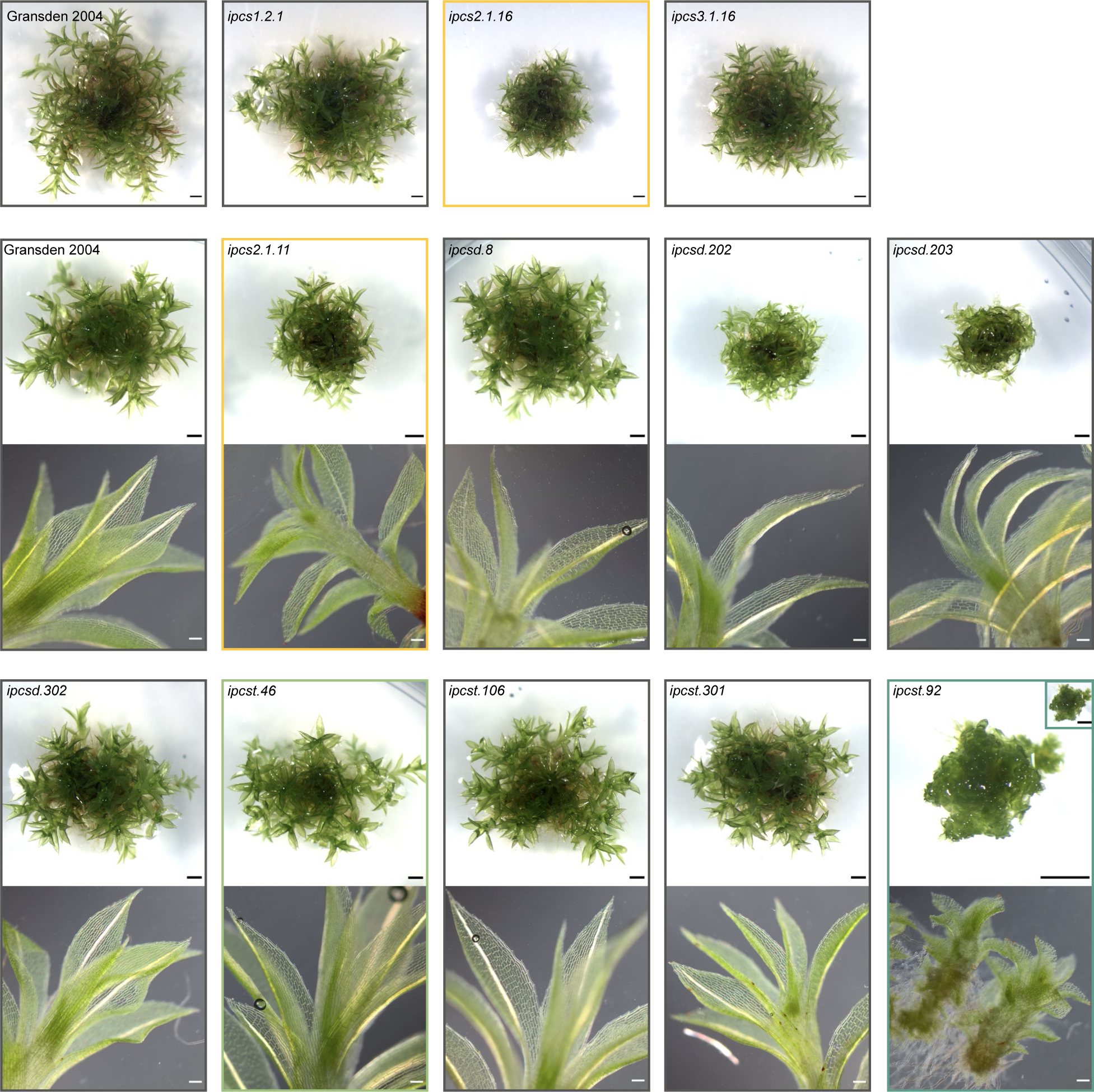
Mature gametophore colonies of *ipcs* single mutant (top row) and selected single and higher-order mutants focus on for analysis. For the selected lines (i.e., all but the top row) colonies of all genotypes were grown on a single plate. Black scale bars in stereomicroscope images represent 1 mm, white scale bars in lower, DIC microscope images represent 0.1 mm. The dwarf *ipcst.92* mutant is imaged at the same magnification as other genotypes in the stereomicroscope image inset for direct comparison to other panels, and magnified in the main stereomicroscope frame.

To generate higher-order mutants (doubles, *d*, and triples, *t*), mutagenesis in wild-type, *ipcs2.1.11*, *ipcs3.1.4*, and *ipcs3.1.16* backgrounds were carried out (Supplemental Table 3). Backgrounds for transformation were selected based on efficiency of the guides used to generate them. Here, the *ipcs1.2* guide had proven extremely efficient, therefore retransforming this guide into genetic backgrounds generated with less efficient guides (i.e. *ipcs2.1* and *ipcs3.1*) was the approach that we expected to require the least screening by sequencing downstream. When screening for mutants with new lesions in multiple loci, we first sequenced the genes targeted with the least effective sgRNAs, followed by sequencing within this population at the sites targeted by most effective guides. This reduced the frequency of sequencing wild-type loci.

We never recovered a triple knock-out, or even an *ipcs1 ipcs2* double null, consistent with the notion that full loss of function would be lethal. Most knock-down higher-order mutants were indistinguishable from the wild type, though mutants including loss-of-function lesions in the *ipcs2* locus (i.e. *ipcsd.202* and *ipcsd.203*) retained the subtle, dwarf phenotype observed in *ipcs2* singles (Figure 2). Exceptionally, one of the triple mutants, *ipcst.92*, exhibited an obvious developmental phenotype. *ipcst.92* has frame-shift lesions in both *IPCS1* and *IPCS3*, and a twelve base pair deletion in *IPCS2* (Supplemental Table 3); this is the closest genotype to a triple knock-out that we obtained.

### The *ipcst.92* mutant phenotype is caused by cumulative deficiency of all three ***IPCS* genes**

To verify that the phenotype of the *ipcst.92* mutant was indeed caused by the three lesions in *IPCS* genes, and not an off-target mutation elsewhere in the genome, we attempted to obtain a second, similarly severe independent allele. We screened hundreds of lines by sequencing, and thousands by phenotype by looking for similarity to *ipcst.92*, in order to recover additional alleles. Unfortunately, we were not successful in isolating an additional mutant allele.

Another strategy to confirm that the phenotype of *ipcst.92* is truly due to *IPCS* gene lesions is to complement the mutation at one of the three *IPCS* gene loci. We replaced the mutant *ipcs2* allele in *ipcst.92* with wild-type *IPCS2* via homologous recombination (HR)-mediated gene targeting. A random selection of recovered lines was genotyped by sequencing the *IPCS2* locus, as well as a subpopulation visually selected based on recovery of the wild-type phenotype (Figure 3A). Lines displaying the *ipcst.92-*like phenotype had retained the *ipcst.92-*derived *ipcs2* lesion, whereas lines with a wild-type-like phenotype contained the wild-type copy of *IPCS2*. Among the wild-type-like lines, 8 were found to have the wild-type copy of *IPCS2* integrated in the genome, at the correct genomic location (Figure 3A).

**Figure 3:**
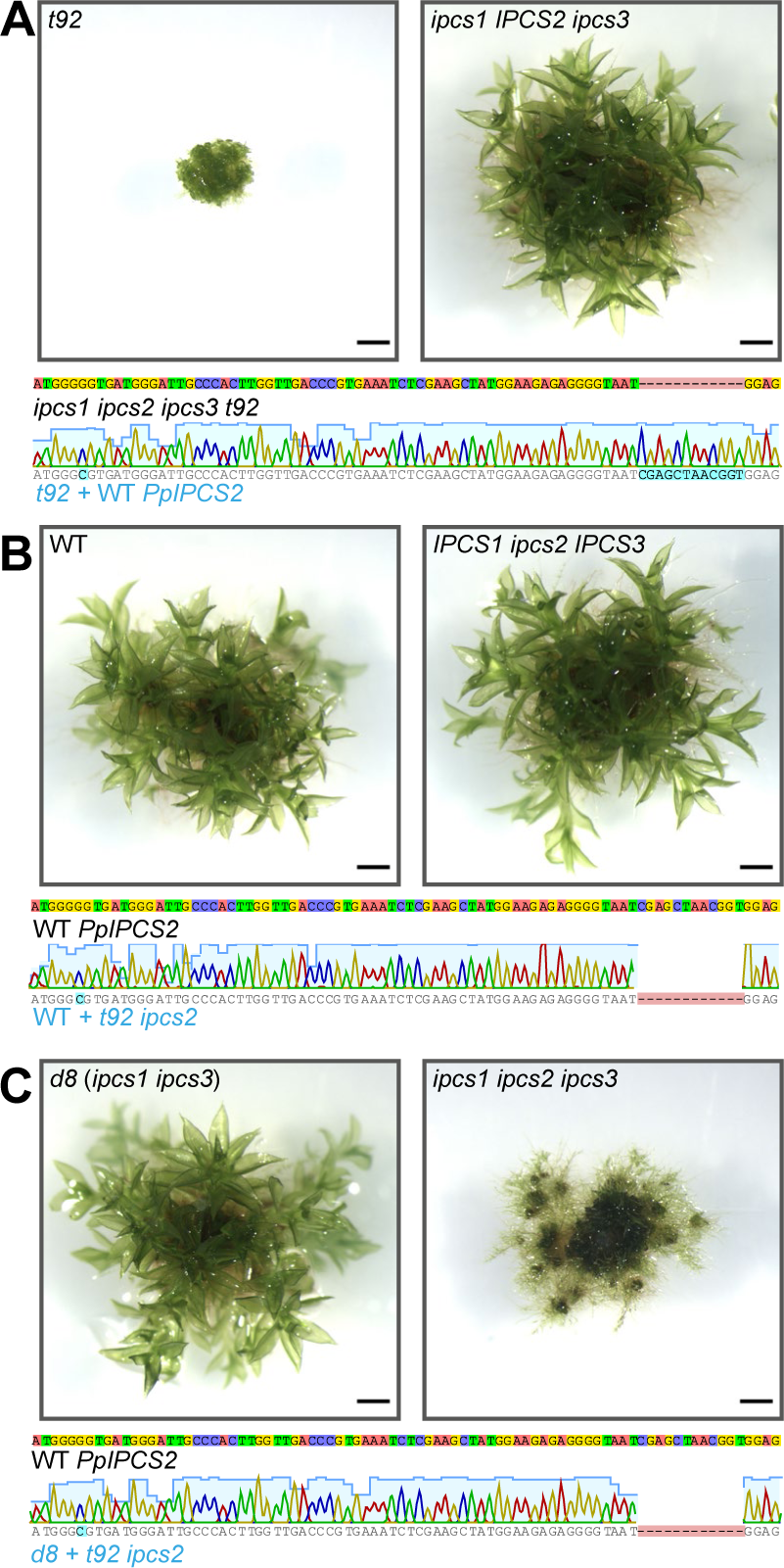
Representative gametophore colonies of **A** The *ipcs1 ipcs2 ipcs3 t92* mutant complemented with a wild-type copy of *IPCS2* cloned from the wild type **B** Gransden 2004 wild type and **C** *ipcs1 ipcs3 d8* transformed with the mutant copy of *ipcs2* cloned from the *ipcs1 ipcs2 ipcs3 t92* mutant. The wild-type *IPCS2* sequence is sufficient to restore a wild-type-like phenotype in the *ipcst.92* triple mutant. The *IPCS1 ipcs2 IPCS3* genotype created by transforming the wild-type background maintains a wild-type like phenotype, while the *ipcs1 ipcs2 ipcs3* line created by transforming *ipcs1 ipcs3* has a similar phenotype to the original *ipcs1 ipcs2 ipcs3 t92* mutant. Colonies of all genotypes were grown on a single plate, and all scale bars represent 1 mm. Sequence reads from the complemented and replication lines are taken from PCR product spanning a region external to the sequence included in the HR sites of the plasmid, ensuring genomic integration of the sequenced read. The silent G>C mutation introduced on HR plasmids is present in all complemented and replicated lines.

A reciprocal experiment was carried out using the mutant copy of *ipcs2* from the *ipcst.92* mutant to replace the wild-type copy of *IPCS2* in either wild type (Figure 3B), or in an *ipcs1 ipcs3* double mutant, *ipcsd.8* (Figure 3C), to replicate the unique *ipcst.92* phenotype. Of all of the recovered lines, only three were observed to have an *ipcst.92*-like phenotype, and this only occurred in the *ipcsd.8* mutant background (Figure 3C). To rule out the possibility that we did not obtain any *ipcst.92*-like lines in the wild-type background by chance, and that these would have presented an *ipcst.92*-like phenotype, we genotyped and sequenced all of the replication lines in the wild-type background. A single line was identified having the mutant *ipcs2* allele from *ipcst.92* integrated in the genome, which clearly maintained a wild-type growth phenotype (Figure 3B).

Altogether, these results indicate that the unique *ipcst.92* phenotype is truly caused by mutations in *IPCS* genes, not off-target lesions. That an *ipcst.92* phenotype was recreated by introducing *ipcs2* from *ipcst.92* in the *ipcsd.8* but not in wild type indicates that a cumulative loss of *ipcs* activity in the *ipcst.92* mutant is responsible for its obvious developmental phenotype, and this phenotype is not a consequence of the specific *ipcs2* lesion in *ipcst.92* (for a technical overview of the experiment, see Supplemental Figure 2).

### Total GIPC content is reduced in the most severe *ipcs* mutants, and total IPC content increases in some mutant alleles

To evaluate the effects of *IPCS* mutations, we measured sphingolipid content with targeted UPLC-nanoESI-MS/MS analysis. Beginning with the single knock-out mutants, we compared our results after normalizing LCMS signals against either FAMEs content determined with an aliquot of each sample, or against sample dry weight. As an example, measurements from protonema of the single mutants are presented in Supplemental Figure 3. Though neither approach is perfect, we reasoned that normalization against FAMEs is a more logical choice: Dry weight is more susceptible to differences in cell size, shape, and cell wall content, whereas FAMEs provided an approximation of the total amount of acyl lipids. Therefore, we only used FAMEs for normalization in subsequent experiments. Unfortunately, absolute quantitation is not possible for GIPCs as appropriate chemical standards are not widely available.

In the single mutants, few substantial, consistent differences were observed in the free ceramides, hexosyl ceramides (HexCers), or GIPCs for the protonema samples taken from single mutants, consistent with the notion that the three *IPCS*s are functionally redundant. There is a reduction in total Hex-GIPC levels in the *ipcs2* single mutants when normalized against dry weight, but this was not significant with FAMEs normalization (Supplemental Figure 3). There was also a substantial increase in free ceramide levels in *ipcs2* mutants, but this was only true when normalized against FAMEs, not dry weight. These weak phenotypes were difficult to trust due to the difference in outcome with different data processing.

We next compared sphingolipid profiles of double and triple mutants, now with FAMEs normalization only. For these mutants, we also used gametophore tissue, not protonema, as we noticed that the greatest phenotypic differences were obvious during this developmental stage when considering the entire mutant collection. The most relevant mutants and controls (Gransden, *ipcs2.1.11*, *ipcst.46*, and *ipcst.92*) are presented in Figure 4, and additional lines are presented in Supplemental Figure 4. We again observed reductions in Hex-GIPC content of *ipcs2.1.11* and higher-order mutants containing a lesion in *IPCS2*, including *ipcsd.202*, *ipcsd.203*, *ipcst.46*, and *ipcst.92*, which affected both A- and B-series and different ceramide backbones. The reductions in *ipcs2.1.11*, *ipcsd.202*, *ipcsd.203*, and *ipcst.46* were subtle and variable between replicates depending on the exact age and treatment of material, while the GIPC reductions in *ipcst.92* were consistently severe, with the mutant accumulating only 15-30 % of the wild-type total Hex-GIPC levels and more severe reductions in Hex-Hex-GIPCs. Notably, reductions were mild or non-existent in HexNAc-GIPCs compared to Hex-GIPCs, with a similar trend between Hex-HexNAc-GIPCs and Hex-Hex-GIPCs (Figure 4A).

**Figure 4:**
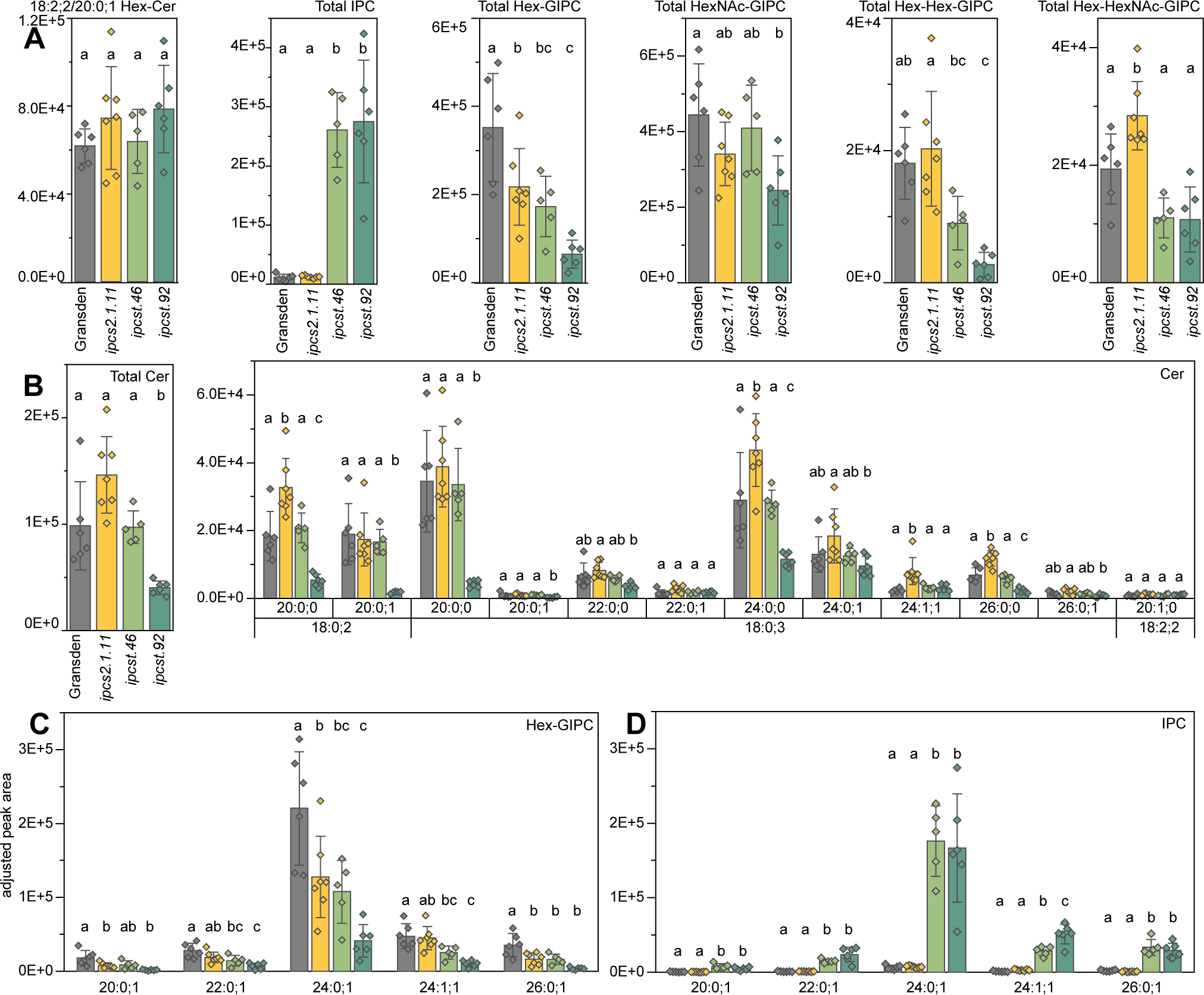
Sphingolipid measurements for selected *ipcs* single and triple mutants. Values are individual MRM peak areas normalized to the total FAME quantity measured for each sample. **A** Total amounts of different classes of sphingolipids. Only the peak area from 18:2;2/20:0;1 HexCer is shown, which accounts for >95% of HexCer in *P. patens*, otherwise the values are the total summed areas for all clearly identified molecular species. **B** Free ceramide total content and composition. **C,** Hex-GIPC composition, and **D**, IPC composition. For B and C, the fatty acid moiety indicated on the y-axis, and the LCB moiety was 18:0;3 for all chemical species. Bars represent averages of five to seven biological replicates, error bars are standard deviation. Letters indicate significance at P < .05 detemined by one-way ANOVA with Tukey’s *post-hoc* test.

Unlike reported *IPCS*-deficient *A. thaliana* lines, there was no increase in free ceramide levels in the most severe *Ppipcs* mutants. The only notable difference we observed was a substantial decrease in accumulation of some specific ceramide species in the *ipcst.92* triple mutant, most obviously 18:0;2/20:0;0, 18:0;2/20:0;1, 18:0;3/20:0;0, and 18:0;3/24:0;0 which resulted in a reduction in the total free ceramide content (Figure 4B). These measurements support the annotations of *IPCS1 IPCS2* and *IPCS3* as true IPCSs, and correlate their weak (*ipcs2*, *ipcsd.202*, *ipcsd.203*, *ipcst.46*) and strong (*ipcst.92*) developmental phenotypes with different degrees of reduction in Hex-GIPC and Hex-Hex-GIPC content (Figure 4A, C).

Most surprisingly, we observed a dramatic increase in IPCs in the *ipcst.46* and *ipcst.92* mutants (Figure 4A, D). This was unexpected because (1) these direct products of the mutagenized *ipcs* gene products were expected to decrease; (2) the quantities of the GIPC end products indeed do decrease; and (3) the *ipcst.46* and *ipcst.92* triple mutants are otherwise dissimilar in their chemotypes and growth phenotypes. We validated the structure of the 18:0;3/24:0;1 IPCs detected in the *ipcst.46* and *ipcst.92* mutants with product ion scans (Figure 5A), and could observe loss of inositol monophosphate (Figure 5B) with positive ionization. The same fragmentation could be observed with the 18:0;3/22:0;1 (Figure 5C) and 18:0;3/24:1;1 (Figure 5D) IPCs. The retention time shifts between these IPCs were consistent with their different fatty acyl desaturations and chain lengths, and their chemical identities were consistent with the Hex-GIPCs that accumulate downstream. Therefore, although the accumulation of IPCs in the mutants is unusual, our data consistently support their correct identification.

**Figure 5:**
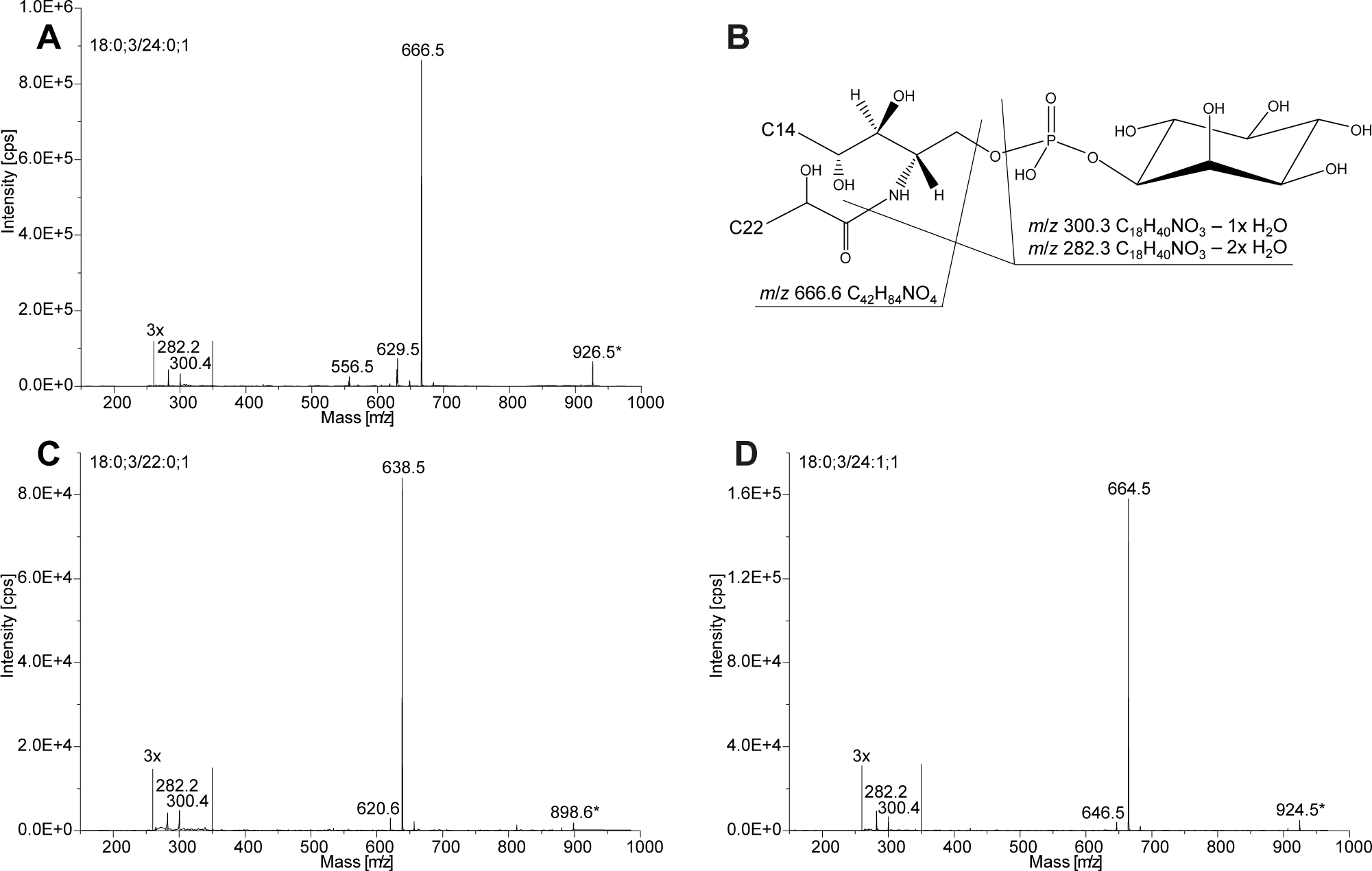
Verification of the inositol phosphorylceramide structures. Fragment ion analysis by UPLC-nanoESI-MS/MS of IPC (A) 18:0;3/24:0;1 (m/z 926.5) in positive ionization mode with a collision energy of 40 eV. (B) Loss of the phosphoinositol-containing headgroup leads to the ceramide fragment of m/z 666.6 and the dehydrated long-chain base fragments m/z 282.2 and 300.4. Fragment ion analysis is also shown for (C)18:0;3/22:0;1 (m/z 898.6), and (D) 18:0;3/24:1;1 (m/z 924.5).

### Accumulation of IPCs in *ipcst.46* and *ipcst.92* is a consequence of their similar mutations in *IPCS2*, not overall loss of *IPCS* activity

The genotypes of the *ipcst.46* and *ipcst.92* mutants are dissimilar. *ipcst.92* has presumed total loss-of-function lesions in *IPCS1* and *IPCS3*, and a partial loss-of-function lesion in *IPCS2*. The *ipcst.46* mutant only has a presumed knock-out lesion in *IPCS3* whereas the lesions in both *IPCS1* and *IPCS2* are presumably only knock down: the *ipcst.46* lesion in *IPCS1* is very likely fully functional, having only a T to NS substitution, and the lesion in *IPCS2* is an 11 amino acid substitution at the same location as that in the *ipcst.92* copy of *IPCS2*, which does maintain sufficient activity to support growth, albeit impaired. The only feature we identified that was common to these two mutant alleles and exclusive from all the others we screened was that they both had short, in-frame deletions near the 5’ end of *IPCS2*. We hypothesized that these specific lesions were responsible for the accumulation of IPCs. We therefore measured sphingolipids in the complementation and replication lines where we had replaced the *IPCS2/ipcs2* loci for confirmation of the causality of the *ipcst.92* phenotype: the singular wild type line with *ipcs2* (i.e. *IPCS1 IPCS2 IPCS3* > *IPCS1 ipcs2^t92^ IPCS3*, simplified wt>t92), three *ipcsd.8* with *ipcs2* (i.e. *ipcs1^d8^ IPCS2 ipcsd ^d8^* > *ipcs1 ^d8^ ipcs2^t92^ ipcs3 ^d8^*, simplified d8>t92), and three *ipcst.92* lines complemented with wild-type *IPCS2* (i.e. *ipcs1 ^t92^ ipcs2 ^t92^ ipcs3 ^t92^* > *ipcs1 ^t92^ IPCS2 ipcs3 ^t92^*, simplified t92>d) (Figure 6, Supplemental Figure 5).

**Figure 6:**
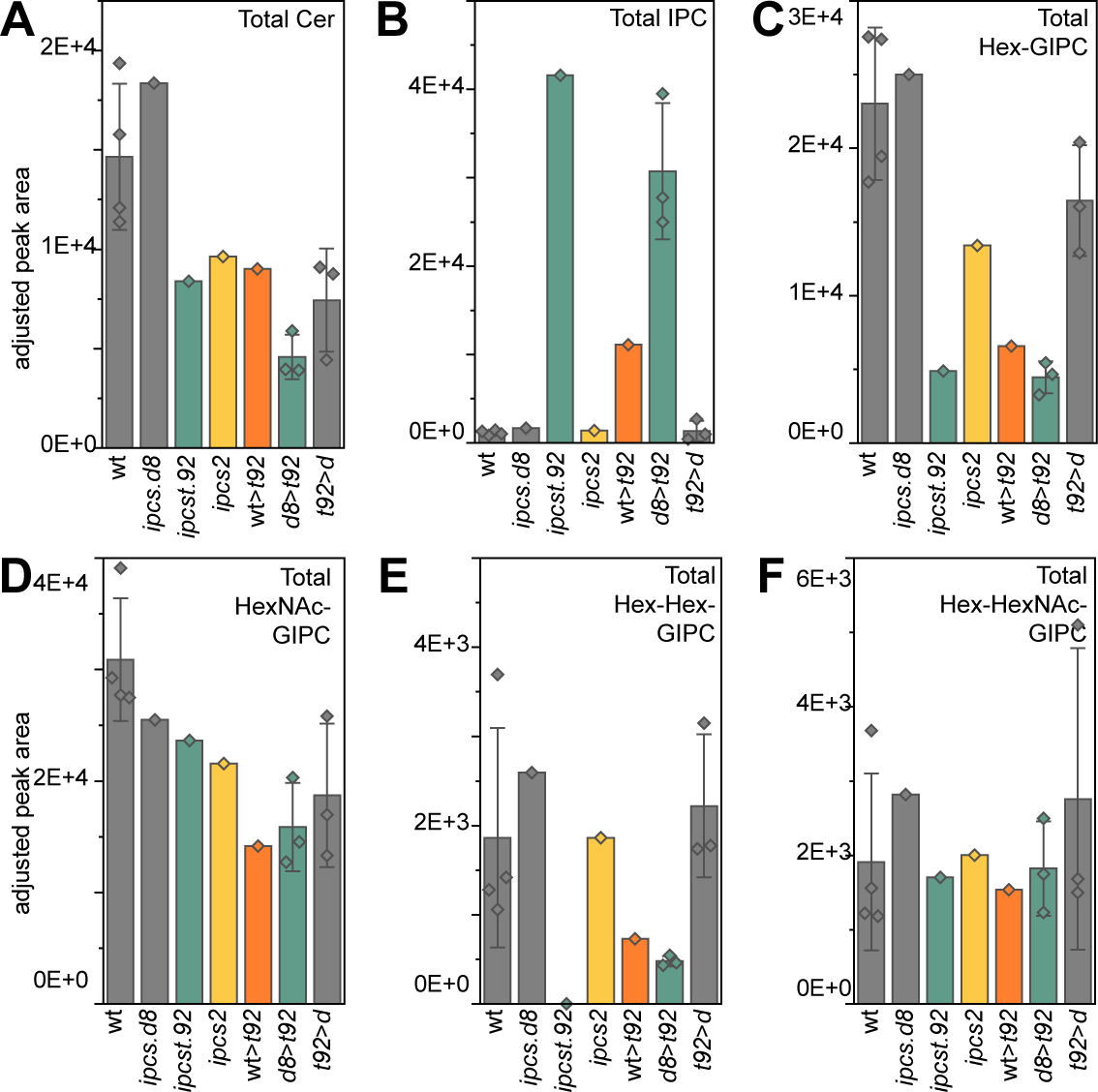
Sphingolipid measurements for background and *ipcst.92* complemented and genotype replicated lines. Values are individual MRM peak areas normalized to the total FAME quantity measured for each sample. **A** Total free ceramides **B** total IPCs **C** total Hex-GIPCs **D** total HexNAc-GIPCs **E** total Hex-Hex-GIPCs **F** total Hex-HexNAc-GIPCs. Bars represent either single samples or averages of three or four biological replicates, where error bars are present they represent standard deviation.

The *ipcst.92* replication lines in the *ipcsd.8* mutant background (d8>t92) had a nearly identical chemotype to the original *ipcst.92* mutant, as expected. The *ipcst.92* complemented lines (t92>d), the *ipcsd.8*, and wild type were indistinguishable. Most importantly, the *ipcst.92* replication line in the wild-type background (wt>t92), having functional copies of *IPCS1* and *IPCS3*, had a chemotype intermediate to the wild type and *ipcst.92*. IPCs accumulated in the wt>t92 line to an extent that was exponential to the levels in the wild-type background, though still less than in the *ipcst.92* triple mutant. This supports our hypothesis that the *ipcs2* lesion is specifically responsible for IPC accumulation.

Perhaps surprisingly, GIPC levels were reduced in wt>t92 to levels similar to *ipcst.92* and the *ipcst.92* replication lines in the *ipcsd.8* background. We expected reduced GIPC levels in wt>t92, however we had anticipated that this deficiency would be similar or weaker than the *ipcs2.1.11* single knock-out mutant, as both off these genotypes have functional *IPCS1* and *IPCS3*. This result suggests that the *ipcst.92* copy of *ipcs2*, while driving the accumulation of IPCs, can produce a more severe GIPC deficiency than entirely losing gene function, perhaps by dis-regulation of the pathway.

### IPCS2 proteins in *ipcst.46* and *ipcst.92* retain their metabolic activity, but their topology may be modified

We can only explain the accumulation of IPCs and simultaneous mild (*ipcst.46*) or severe (*ipcst.92*) reduction in GIPCs in the triple mutants by their expressed, modified IPCS2 enzymes retaining activity in a way that makes their direct products inaccessible for downstream processing. *ipcst.46* would still accumulate GIPCs due to presumed functionality of its *ipcs1* gene product, which has only a weak lesion (Supplemental Table 3). To directly compare the functionality of the IPCS2 proteins of wild type and both triple mutants, we expressed them in the conditional lethal IPCS-deficient yeast strain *YPH499-HIS3-GAL1_pro_:AUR1*. All three IPCS2 proteins could rescue growth of the mutant under restrictive conditions (Figure 7A, B). We then extracted total lipids for sphingolipid measurements (Figure 7C-H, Supplemental Figure 6). With wild-type *PpIPCS2* and *Ppipcs2* from the *t46* mutant, there was an overall increase in mannose inositol phosphorylceramide (MIPC) accumulation, and enrichment in both IPCs and MIPCs containing an 18:0;3 LCB moiety. The preference for 18:0;3 LCBs fits with the expected substrate preference for these *P. patens* enzymes. With the *Ppipcs2* sequence from the *t92* mutant, although there was complementation and accumulation of IPCs, MIPCs, and mannosyl di-inositol phosphorylceramides (M(IP)_2_Cs), the overall levels of IPCs and MIPCs were reduced compared to the background strain as well as the wild type- and *t46*-complemented lines. These results indicate that all three *PpIPCS2/Ppipcs2* sequences produce functional IPCS enzymes in yeast, but that *ipcs2* from *t92* has reduced functionality. Most importantly, whatever the reason for increased accumulation of IPCs in the *ipcst.46* and *ipcst.92* mutant plants, this effect was not reproduced in yeast cells.

**Figure 7:**
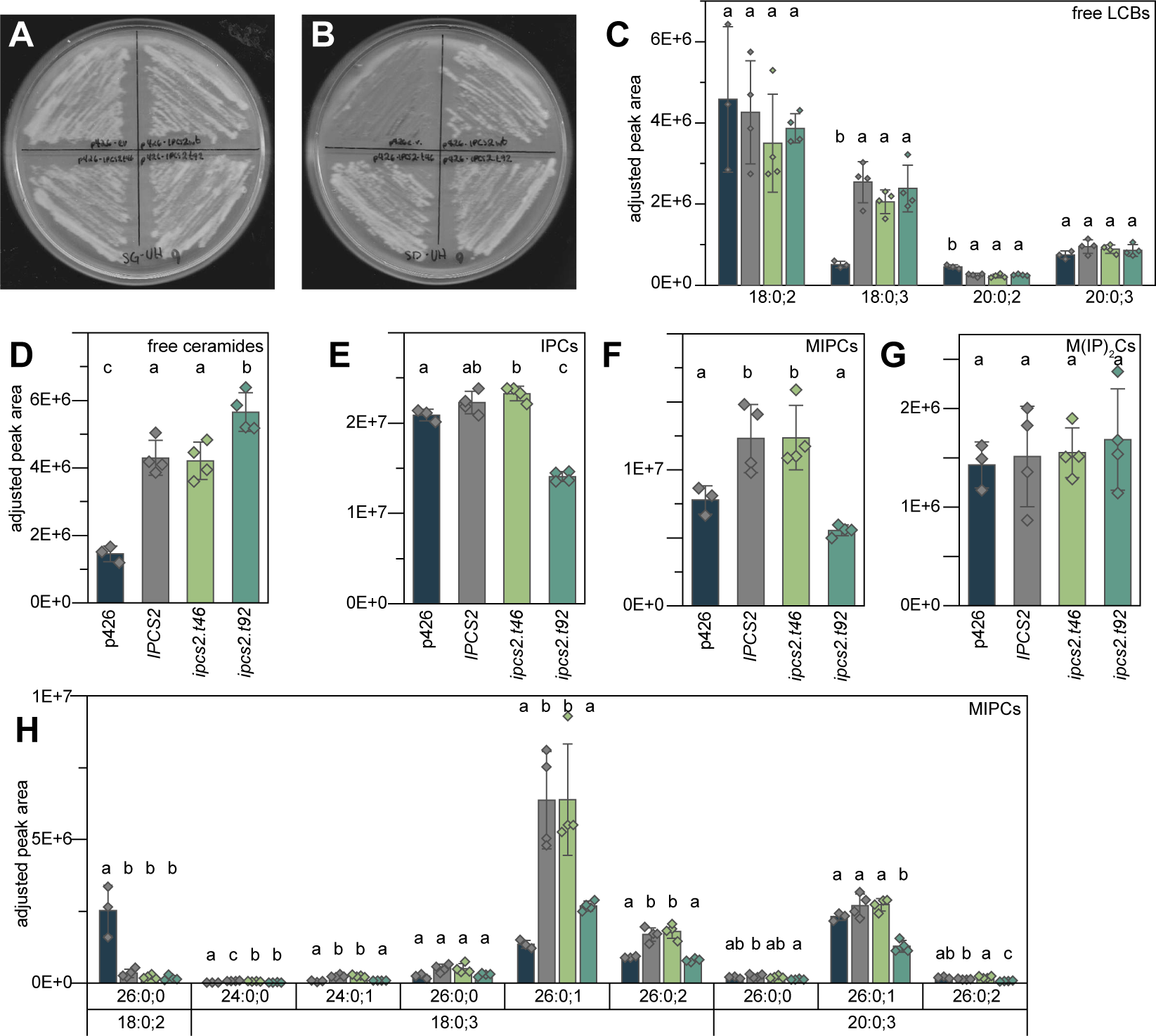
Complementation of *S. cerevisiae* YPH499-HIS-GAL1::AUR1 conditional mutant with *P. patens IPCS2* wild type, and mutant *ipcs2* alleles cloned from *ipcst.46* and *ipcst.92*. **A** Control cells grown on galactose-containing medium (SG-UH) express endogenous *AUR1* and survive independent of transgene expression. **B** *AUR1* expression is surpressed on glucose-containing medium (SD-UH). Cells with only the empty vector (p426e) are not viable, but cells expressing the wild-type, *ipcst.46*, or *ipcst.92* copies of *P. patens IPCS2* are complemented. **C-H** Sphingolipid measurements for *S. cerevisiae* YPH499-HIS-GAL1::AUR1 with the empty vector cultivated on SG-UH (three replicates), and *S. cerevisiae* YPH499-HIS-GAL1:: AUR1 with *P. patens IPCS2* gene constructs grown on SD-UH (four replicates each). Bars represent averages with standard deviation. Letters indicate significance at *P* < .05 detemined by one-way ANOVA with Tukey’s *post-hoc* test.

Wild-type and *ipcst.92* IPCS2 with C-terminal fusions to eYFP showed similar localization patterns when transiently expressed in *Nicotiana benthamiana* leaf, indicating that the IPCS gene products are not mis-localized in any drastic, obvious way (Supplemental Figure 7).

The IPCS2 proteins in *ipcst.46* and *ipcst.92* both contain short deletions, eleven and four amino acids each, respectively (Figure 8A). Structural predictions generated with Alphafold, and topology predictions generated with TMHMM, CCTOP, and TOPCONs, place the lesion site of both alleles in an N-terminal region that precedes the first transmembrane domain (TMD) (Figure 8B) opposite to the lumenal catalytic site. Notably, in *ipcst.46*, the deletion removes three of four positively-charged residues that precede the first TMD. We expect that removal of this positive charge will impact cotranslational insertion of the peptide into the ER, as these would otherwise interact with the translocation machinery to retain the N-terminus on the cytosolic side of the membrane (Rapoport et al., 2004). In the absence of positive charge, the N-terminus can be released into the ER lumen, inverting the topology of the first and all downstream TMDs. Although the four-residue deletion of *ipcst.92* does not remove any positive charges, it does introduce a new translational start site (ATG) downstream of all four positive residues, which could produce an alternative peptide with inverted topology. IPCS2 protein inversions in both mutants could nicely explain their accumulation of IPCs: when the catalytic site is flipped to face the cytosol, IPC products would be released in the cytosolic leaflet of the membrane, where they would be inaccessible to the enzymes responsible for glycosylations to produce mature GIPCs (Figure 8C, D).

**Figure 8:**
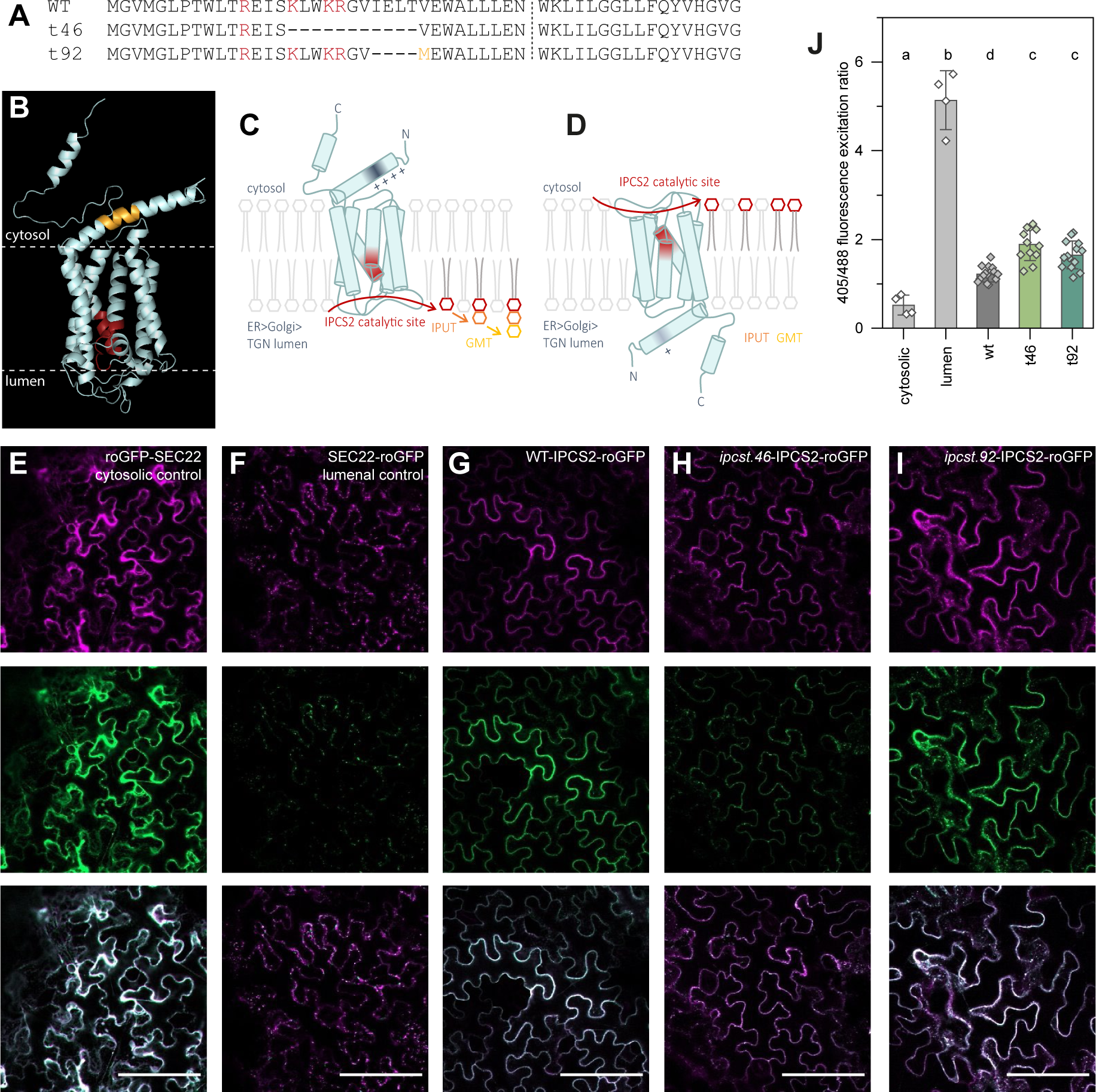
Functional characterization of IPCS2 proteins from *ipcst.46* and *ipcst.92*. **A** Sequence alignment of wild type and mutant IPCS2 proteins. Positively charged residues are highlighted red, and a new methionine residue, or translational start, introduced by the out-of-frame 12 bp deletion in *ipcst.92* is highlighted gold. The beginning of the first predicted transmembrane domain is marked with a dashed line. **B** Structural model of wild type IPCS2. Model was generated with alphafold, and the overlaid topology prediction with TMHMM. Conserved residues associated with catalysis are highlighted red, and the sequence targeted for restriction by Cas9 is highlighted gold. **C**&**D**: Models of (**C**) wild type, and (**D**) *ipcst.46* mutant IPCS2 protein topology and hypothesised mis-localisation of IPC intermediates in the cytosolic leaflet of the endomembrane. **E-I** Representative merged images of roGFP2 fusions excited with both 488 and 405 nm light. Emission from excitation with the 488 nm laser is false-coloured green, and emission from excitation with the 405 nm laser is false-coloured magenta. White values of the display curve were reduced for individual images to enhance brightness, as fluorescence intensity was lower for IPCS2-roGFP2 fusions than for controls; the white value was kept identical for the two channels within individual images. **J** 405 nm/488 nm fluorescence excitation ratios. Individual data points are ratios calculated from the raw arithmetic mean intensity of the two channels in individual images, bars represent averages with standard deviation. Letters indicate significance at P < .01 detemined by one-way ANOVA with Tukey’s *post-hoc* test.

We tested this model by cloning wild type and mutant IPCS2 proteins with C-terminal fusions to redox-sensitive GFP (roGFP2). roGFP2 can be used as a topology sensor due to the strong glutathione redox difference between the cytosol (Figure 8E) and endomembrane lumen (Brach et al., 2009) (Figure 8F). When transiently expressed in *N. benthamiana* leaf, the wild-type IPCS2-roGFP2 fusion showed a cytosolic roGFP2 fluorescence pattern, supporting topology predictions. By comparison, IPCS2 from *ipcst.46* and *ipcst.92* were more oxidized, suggesting that some of the roGFP2 tag could be localized to the endomembrane lumen when fused to these mutant alleles of IPCS2. However, the difference we observed between the wild-type and mutants was substantially weaker than for the control endomembrane proteins. Therefore, this result and our model are not absolutely conclusive and are open to debate. Nevertheless, misfolding and modified topology, for a fraction of the expressed IPCS2 protein in the two triple mutants, remains the most logical explanation we have that fits both the chemotypes and the specific mutant lesions.

## Discussion

Methods for genome editing of *P. patens* with CRISPR/Cas have developed rapidly in recent years, including tools for knock-out mutagenesis, base editing, and gene targeting. These, paired with the simple developmental patterning and consequent ease of manipulation and observation of *P. patens*, have made it an exciting model that has expanded in popularity. Here, we’ve made simple adaptations to available approaches designed for knock-out mutagenesis to specifically screen for knock-down mutants. In this way, we can better understand essential genes and gene families, and the fundamental biological processes that their products contribute to. Our approach offers different advantages from recent protocols for base editing and prime editing in *P. patens* (Guyon-Debast et al., 2021; Perroud et al., 2023), in that it deliberately produces populations of diverse alleles. This allows for variation in allele severity, and increases the chances of finding something unexpected. Our approach is only possible because *P. patens* protoplasts are easily transformed and, most critically, totipotent. This is the key factor that enables collection of large numbers of distinct mutant alleles.

We were successful in isolating diverse alleles of *ipcs1-3* with a gradient of sphingolipid deficiencies, including up to an 85 % reduction in Hex-GIPCs in gametophores of *ipcst.92*. In *A. thaliana*, the most severe *ipcs* mutant alleles are non-viable, and the strongest IPCS-deficient lines that are tractable display only modest reductions in GIPCs (Wang et al., 2008; Mina et al., 2010; Ito et al., 2021). Therefore, the *P. patens* mutants offer a unique opportunity to observe the effects of GIPCs on membrane dynamics and development. The GIPC-deficient *ipcst.92* is stunted and displays obvious developmental phenotypes. It is remarkably lucky that in this mutant we can link these phenotypes directly to GIPCs; in most *ipcs* mutants GIPC depletion is paired with ceramide substrate accumulation, and the effects of this potent PCD signal are impossible to distinguish from GIPC phenotypes. We infer that free ceramides do not accumulate in *ipcst.92* because they are still processed to IPCs. We can rule out the possibility that IPC accumulation causes growth defects because of the wild-type-like phenotypes of *ipcst.46*, and the *ipcst.92* replication line in wild-type background (wt>t92, Figures 3 and 5).

Accumulation of IPCs in *ipcst.46* and *ipcst.92* was an exciting and unanticipated finding, that we can only explain by mis-localization or folding of IPCS2 protein. Based on the *ipcs2* lesions in these two alleles, inverted topology of the IPCS2 protein products and consequent mis-localization of IPC metabolic intermediates is the most logical explanation, which is supported, albeit weakly, by the redox-based topology analysis. We expect that further investigation of these mutants could provide insight into trafficking and eventual localization of GIPCs in the apoplastic PM leaflet.

## Methods

### Candidate gene identification

Candidate *IPCS* genes were identified by a BLAST search of the *Physcomitrium patens* ecotype Gransden V.3 genome (Lang et al., 2018) for homologs of characterized *IPCS* genes from *Arabidopsis thaliana* via the Phytozome server (Goodstein et al., 2012). Gene loci were named: *PpIPCS1* (Pp3c8_19970V3.1, LOC112286074), *PpIPCS2* (Pp3c23_1080V3.1, LOC112275922), and *PpIPCS3* (Pp3c20_10120V3.1, LOC112272995). Peptide sequences of the gene products were aligned in MUSCLE (Supplemental Figure 1) and percent identities computed by Clustal 2.1(Madeira et al., 2022) (Figure 1B). Sequence analysis, construct planning, and verification was carried out using Geneious 8.1.8. (http://www.geneious.com) and A plasmid Editor (ApE) (https://jorgensen.biology.utah.edu/wayned/ape/)(Davis and Jorgensen, 2022).

### Cultivation of Physcomitrium patens

*Physcomitrium patens* ecotype Gransden 2004 was obtained from the International Moss Stock Center (IMSC; https://www.moss-stock-center.org/en/), strain 40001. Plants were maintained at 25 °C under long-day conditions (16 h light/8 h dark) with 105-120 μmol m^-2^ s^-1^ radiation. Filamentous protonema tissue was cultivated on BCD-AT medium (1 mM MgSO_4_, 1.84 mM KH_2_PO_4_, 10 mM KNO_3_, 45 μM FeSO_4_, 5 mM ammonium tartrate, 1 mM CaCl_2_, Hoagland’s trace elements, 0.55 % plant agar) overlaid with sterile cellophane discs, and the cultures were propagated by disruption in sterile tap water with an IKA ULTRA-TURRAX homogenizer every 1-2 weeks. Development of gametophores was induced by transferring small pieces of tissue to BCD medium (BCD-AT lacking ammonium tartrate). Gametophores were sub-cultivated every 4-6 weeks (Maronova and Kalyna, 2016).

### RNA extraction, RNA sequencing, and cDNA synthesis

RNA was extracted from mature, lyophilized gametophores of wild-type Gransden and the *ipcst.46* mutant. 5 mg lyophilized material was used for each of three independently grown and harvested replicates per genotype. Total RNA was extracted with the Omega Bio-Tek E.Z.N.A. Plant RNA Kit, quantified, and 1 μg of total RNA was treated with DNaseI (Thermo Fisher Scientific). RNA quality was controlled, an Illumina_TruSeq mRNA-SeqSample library was prepared, RNA sequencing was performed, and data was analyzed by the University of Göttingen Next Generation Sequencing Integrative Genomics Core Unit.

cDNA produced for cloning was synthesized from DNase-treated RNA, with RevertAid H Minus Reverse Transcriptase (Thermo Fisher Scientific), according to the manufacturer’s instructions.

### Complementation of the conditional Saccharomyces cerevisiae YPH499-HIS3-GAL1_pro_:AUR1 mutant

Coding sequences of *PpIPCS1*, *PpIPCS2*, and *PpIPCS3* were amplified from either cDNA or from genomic DNA in the case of sequences lacking introns, prepared from protonema and gametophore material of wild-type Gransden 2004 *Physcomitrium patens*. The *AtIPCS2* coding sequence was prepared from *Arabidopsis thaliana* cDNA, for use as a positive control (Mina et al., 2010). Primers added a Kozak consensus sequence for optimal translation in *Saccharomyces cerevisiae*, AAAA, directly upstream of the ATG start codon (Nakagawa et al., 2008). All primers for this study are listed in Supplemental Table 4. Coding sequences were ligated into the p426 expression vector by conventional restriction enzyme cloning, and plasmid sequences were verified by restriction digests and Sanger sequencing. Expression plasmids were transformed in parallel into aliquots of *YPH499-HIS3-GAL1_pro_:AUR1* cells via the LiAc/SS carrier DNA/PEG method(Gietz and Schiestl, 2007). To maintain the mutant strain and select for transgenic cells, yeast was cultivated in or on synthetic complete dropout medium lacking the appropriate nutrients and carbohydrate supplementation.

### Stereomicroscope and DIC imaging of *Physcomitium patens*

After ventilating plates under a sterile bench to reduce humidity and condensation, mutant and wild-type gametophores were photographed directly on covered petri dishes, using a Retiga R6 camera on an Olympus SZX12 stereomicroscope. DIC images were captured with a Zeiss Axiocam 208 color camera with 5 X magnification.

### CRISPR/Cas9 mutagenesis

The pBNRF selection plasmid and Act1_pro_-Cas9 nuclease plasmid were kindly provided by Prof. Fabien Nogué (INRA Versailles), and used following protocols established in their lab (Collonnier et al., 2017; Lopez-Obando et al., 2016). For single guide cloning, a custom vector (pUCRISPR) was generated in the pUC57 vector backbone, using Genscript’s gene synthesis service, modelled on synthetic genes described (Lopez-Obando et al., 2016). The insert consisted of the *Physcomitrium patens* U6 promoter sequence, followed by dual BbsI restriction sites for guide cloning, followed by the optimized *tracrRNA* sequence described (Castel et al., 2019) (Supplemental Figure 8). Single guide sequences were selected using the CRISPOR platform (http://crispor.tefor.net/) (Concordet and Haeussler, 2018). Guides were ordered as single-stranded synthetic oligonucleotides, phosphorylated with Phosphonucleotide Kinase (PNK), annealed into duplexes, and ligated into BbsI-cut and dephosphorylated pUCRISPR vector. Plasmids were verified by Sanger sequencing. Following transformation (below), individual mutants were identified by amplification of an approximately 0.5 kb region surrounding the target site and Sanger sequencing of the fragment. Although we selected target sites with the fewest likely off-targets and highest predicted efficiency, some of our guides (e.g. *ipcs1.1*, *ipcs2.2*) were less efficient than other guides we used in parallel. The reasons for this are not clear.

### Generation and transformation of *Physcomitrium patens* protoplasts

Protoplast transformation was based on the methods described (Liu and Vidali, 2011; Maronova and Kalyna, 2016; Lopez-Obando et al., 2016). Briefly, protoplasts were prepared by digesting 5-6-day-old protonema from a single 90 mm Petri dish culture, in a 1 % Driselase, 8 % mannitol solution for approximately 3 h. This amount of tissue was sufficient for four parallel transformations. The digested suspension was filtered through a 70 μm sterile filter, and the cells were centrifuged at low speed and washed twice in 8 % mannitol to remove the Driselase. The clean, pelleted protoplasts were re-suspended in 2.5 mL mannitol magnesium solution (0.4 M mannitol, 15 mL MgCl_2_, 4 mM MES pH 5.7), and incubated for 20 min at RT. 600 μL aliquots of the protoplast suspension were then added to prepared mixtures of the required plasmids, which included 10 μg pAct1-Cas9 and equimolar amounts of pBNRF and each pUCRISPR plasmid, for a final amount of plasmid not exceeding 60 μg. 700 μL PEG/Ca solution (4 g PEG4000, 3 mL H_2_O, 2.5 mL 0.8 M mannitol, 1 mL 1 M CaCl_2_) was added to each transformation, gently mixed by inverting, and incubated at RT for 30 min. Afterwards, the solution was diluted with 3 mL W5 solution (154 mM NaCl, 125 CaCl_2_, 5 mM KCl, 2 mM MES pH 5.7), spun down at low speed, and the cells resuspended in 5 mL molten top protoplast recovery medium (PRM-T) (BCD-AT with 10 mM CaCl_2_, 8 % mannitol, 0.4 % plant agar, kept molten in a 42 °C hot water bath). The suspension was split and plated evenly on two bottom protoplast recovery media (PRM-B) (BCD-AT with 10 mM CaCl_2_, 8 % mannitol) plates overlaid with cellophane. After five days of recovery, the cellophane discs carrying the PRM-T-embedded protoplasts were transferred to BCD-AT medium with G418 to select for transformed colonies.

### *ipcst.92* mutant rescue and replication

HR cassettes were generated as synthetic genes by BioCat (BioCat GmbH, Heidelberg, Germany), and directly obtained in the pUC57 vector backbone. The efficiency of HR can be improved by combined use with CRISPR/Cas9 to cleave the targeted genomic region; therefore, we prepared sgRNA constructs that would target sites neighbouring the lesion present in the *ipcst.92* copy of *IPCS2*, and internal to 750 bp regions at either end of the *ipcs2* sequence designated here as the HR target sites. For the wild-type *IPCS2* HR plasmid, the wild-type-like sequence was synthesized with two silent point mutations at the exact sites we targeted with CRISPR/Cas9, so that Cas9 would cleave the *ipcst.92* genomic copy of *IPCS2*, but not the wild-type-like copy present on the plasmid. The mutations maintained the same amino acid sequence using codons that are more frequently used in *P. patens* than the original sequence. Sequences of the expression cassettes in the synthesized plasmids were confirmed by Sanger sequencing. sgRNA plasmids for targeting sequences flanking the region of interest were designed and cloned as described under CRISPR/Cas9 mutagenesis. To generate complementation lines, the HR, both sgRNA, Cas9, and pBNRF plasmids were co-transformed into *ipcst.92* mutant protoplasts. To replicate the *ipcst.92* mutant genotype, the gene copy cloned for HR (here *ipcs2* from *ipcst.92*) was also synthesized so that it included silent point mutations that rendered it immune to CRISPR/Cas9 mutagenesis. HR, both sgRNA, Cas9, and BNRF plasmids were all co-transformed for simultaneous CRISPR/Cas9 mutagenesis and HR. Individual putative rescue and replication colonies were screened by phenotype and subsequently genotyped by sequencing a region that spanned beyond the HR sequence through the *ipcs2* lesion. This was accomplished by amplification and sequencing from a region outside of that cloned for HR, and finding that this sequence is contiguous with the wild-type-like *IPCS2* sequence, including a silent base substitution introduced on the plasmid DNA. In lines with a wild-type appearance where the mutant copy of the gene was sequenced with this approach, we assume that the wild-type copy of the gene cloned for HR was maintained as, and expressed from, episomal DNA.

To screen the entire replication line population in the wild-type background, we amplified a fragment of *IPCS2* using a forward primer specific to the region upstream of the cloned HR sequence, and a reverse primer that anneals to the 16 bp missing in the *ipcs2* mutation of *ipcst.92*. This enabled high-throughput screening of the *ipcst.92* mutation without sequencing. Individuals missing this PCR product were then verified by sequencing the mutagenized locus.

## Sphingolipid Analysis

### Microsome enrichment from *Physcomitrium* protonema and gametophore tissues

Microsomes were prepared from 5-20 mg lyophilized protonema or gametophores, with a consistent dry mass of tissue used within experiments. Lyophilized tissues were ground with a laboratory mixer mill MM 400 (Retsch GmbH, Haan, Germany) using stainless steel beads. The microsome enrichment followed established methods (Abas and Luschnig, 2010), with minor modifications. Briefly, the pulverized tissue was suspended in a fractionation buffer consisting of 150 mM Tris-HCl (pH7.5), 37.5 % sucrose, 7.5 % glycerol, 15 mM EDTA, 15 mM EGTA, and 7.5 mM KCl. After suspension, all steps were performed on ice or at 4 °C. The suspensions were centrifuged at low speed, 1,500 g for 3 min, to remove cell debris, and the supernatants retained. The pellets were re-suspended in a similar fractionation buffer to the first, with 0.75 X concentrations of all components, and the separation repeated. All steps were repeated a third time with a buffer at 0.67 X concentration. All of the supernatants were pooled and diluted 1:1 with water, and centrifuged at 20,817 g for 2 h to pellet the microsomes. The pellets were washed with water and re-spun, then stored at 80 °C until further processing.

### Lipid extractions from *Physcomitrium patens-*derived microsomes, and lyophilized *Saccharomyces cerevisiae* cells

Lipid extraction and analysis was based on published methods (Herrfurth et al., 2021). Briefly, either microsomes or total lyophilized, pulverized material was suspended in 6 mL extraction buffer consisting of isopropanol:hexane:water (60:26:14, v/v/v), vortexed, sonicated, and shaken at 60 °C to fully solubilize lipids. Cell debris was spun down at 800 g for 20 min at room temperature, the supernatant transferred to a fresh tube, and dried under a nitrogen stream. The lipid residue was finally re-suspended in 800 μL extraction buffer for storage under argon gas at -80 °C, or further processed directly.

### Fatty acid methyl ester (FAMEs) derivatization and GC-FID analysis

A 20 % aliquot of the total extract volume was taken for methyl esterification by sulfuric acid-catalyzed methanolysis. The aliquots were dried under a nitrogen stream and re-suspended in 1 mL sulfuric acid solution, and 5 μg 15:0 triacylglycerol standard added to each sample. The derivatization was carried out in an 80 °C water bath for 1 h. The reaction was stopped and FAMEs were extracted by addition of 200 μL 5 M aqueous NaCl solution and 200 μL hexanes. The upper hexane phase was transferred to a clean tube, evaporated under nitrogen, and finally re-suspended in 20 μL acetonitrile for injection into a GC-FID for analysis and total fatty acid quantification. The total fatty acid content was used to normalize LCMS MRM peak areas of individual lipids for relative comparisons.

### Methylamine treatment of lipid extracts

50 % of the total lipid extracts were treated with methylamine to hydrolyze glycerolipids. This provided cleaner background and removed signals that could otherwise be mis-interpreted as false positives in our sphingolipid analysis. This followed a protocol (Herrfurth et al., 2021) adapted from previous work (Markham and Jaworski, 2007). Briefly, the samples were dried down under a nitrogen stream, and the residue re-suspended in 700 μL 33 % (v/v) methylamine in ethanol, and 300 μL H_2_O. The derivatization took place at 50 °C for 1 h, and the solvents subsequently evaporated off under a nitrogen stream. The residue was resuspended in tetrahydrofuran:methanol:water (4:4:1, v/v/v) for injection into the LCMS for analysis.

### UPLC-nanoESI-MS/MS analysis of molecular lipid species

Sphingolipids were separated after methylamine treatment by an ACQUITY UPLC system (Waters Corp., Milford, MA, USA) equipped with an HSS T3 silica-based reversed-phase C18 column (100 mm x 1 mm, 1.8 µL; Waters Corp.), ionized by a chip-based nano-electrospray using TriVersa Nanomate (Advion BioScience, Ithaca, NY, USA) with 5 µm internal diameter nozzles and analyzed by an AB Sciex 6500 QTRAP tandem mass spectrometer (AB Sciex, Framingham, MA, USA) (Herrfurth et al., 2021). The instrument was operated in positive ionization mode of the nanomate and in multiple reaction monitoring (MRM) mode of the mass spectrometer.

Sample injection of 2 µL was performed by an autosampler set at 18 °C and sample separation was conducted at a flow rate of 0.1 mL/min. The solvent system was composed of methanol:20 mM ammonium acetate (3:7, v/v) with 0.1 % acetic acid (v/v) (solvent A) and tetrahydrofuran:methanol:20 mM ammonium acetate (6:3:1, v/v/v) with 0.1 % acetic acid (v/v) (solvent B). For sphingolipid separation except long-chain bases, a linear gradient was applied: start from 65 % B for 2 min; increase to 100 % B in 8 min; hold for 2 min and re-equilibrate to the initial conditions in 4 min. Starting condition of 40 % solvent B was utilized for LCBs. For the MRM detection, precursor ions were [M+H]^+^ and product ions were dehydrated LCB fragments for LCBs, ceramides, and glycosylceramides. The loss of phosphoinositol-containing head groups was used for detection of glycosyl inositol phosphorylceramides.

For structure verification of the inositol phosphorylceramides, product ion scans were performed in the positive ion mode with the precursors [M+H]^+^ and product ion detection was performed in a mass range of 150 to 1000 Da. The collision energy was 40 eV.

### Topology predictions and structural modelling

Topology of PpIPCS2 was predicted with TMHMM (Hallgren et al., 2022), TOPCONS (Tsirigos et al., 2015), and CCTOP (Dobson et al., 2015). The tertiary structure of PpIPCS2 was modelled using Alphafold v2.3.2 (Jumper et al., 2021).

### Topology sensing with roGFP2 in *N. benthamiana*

Our approach followed a method developed and reported previously (Brach et al., 2009). The *IPCS2* coding sequence was amplified from wild type, *ipcst.46*, and *ipcst.92* genomic DNA (*PpIPCS2* contains no introns), and ligated into a pEntry plasmid derived from pUC18, which includes the 35S promoter and a C-terminal fusion to the eYFP fluorophore. Subsequently, eYFP was excised and replaced with the roGFP2 fluorophore, amplified from an expression construct kindly shared by Prof. Stefanie Müller-Schüssele and Julian Ingelfinger (Division of Molecular Botany, Technical University Kaiserslautern). All primers used for assembling these constructs are listed in Supplemental Table S1. The expression cassettes were moved into pCAMBIA binary expression vectors using standard gateway cloning. Cytosolic and lumenal controls are pCM01-35S::roGFP2-SEC22 and pSS01-35S::SEC22-roGFP2 (Brach et al., 2009).

The binary expression vectors were transformed into chemically competent GV3101 *A. tumefaciens* containing the pMP90 Ti plasmid. 3-5 week old *N. benthamiana* leaves were co-infiltrated with the pCAMBIA-35S_pro_::IPCS2-roGFP2 plasmids and pBin61-p19 for expression, and suppression of gene silencing (Qiu et al., 2002). Confocal imaging of abaxial pavement cells was carried out 1-2 days post-infiltration on a Zeiss Axio Imager.Z2, with a Plan-Apochromat 20X/0.8 M27 objective. Scans with the 405 and 488 nm lasers were carried out with frame switching, and the detection window for both channels was 507-534 nm. The 488 nm laser intensity was 0.3 %, the 405 nm laser intensity was 1.2 %, and the detector gain for both was set to 800 V. Images were processed in ZEN 3.2 and ImageJ 1.53f5.1. The 405/488 excitation ratio was calculated from arithmetic mean intensities of all individual images from plants transformed in parallel.

## Acknowledgements

We are grateful for excellent technical support from Sabine Freitag. We thank Prof. Fabien Nogué for generously sharing plasmids and invaluable advice related to CRISPR/Cas9 mutagenesis in *P. patens*. We thank Prof. Ralph T. Schwarz and Dr. med vet. Hosam Shams-Eldin for kindly providing the *YPH499-HIS3-GAL1_pro_:AUR1* yeast strain. Sincere thanks to Prof. Stefanie Müller-Schüssele and Prof. Andreas Meyer for roGFP2 plasmids, and additionally to Prof. Meyer for lengthy discussions and insight related to roGFP2 topology assays. We thank Dr. Patricia Scholz for providing the R script used for statistical analysis. We thank Prof. Jan de Vries and Dr. Tatyana Darienko for access to and kind assistance using their DIC microscope. IF acknowledges funding through the German Research Foundation (DFG: INST 186/822-1, INST 186/1167-1). TMH was supported by a Humboldt Research Fellowship for Postdoctoral Researchers from the Alexander von Humboldt Foundation (CAN 1210075 HFST-P), and a Marie Skłodowska Curie Independent Fellowship (MSCA-IF-EF-ST:892532—SMFP) from the European Research Council.

